# More Structure, Less Accuracy: ESM3’s Binding Prediction Paradox

**DOI:** 10.1101/2024.12.09.627585

**Authors:** Thomas Loux, Dianzhuo Wang, Eugene I. Shakhnovich

## Abstract

This paper investigates the impact of incorporating structural information into the protein-protein interaction predictions made by ESM3, a multimodal protein language model (pLM). We utilized various structural variants as inputs and compared three widely used structure acquisition pipelines—EvoEF2, Gromacs, and Rosetta Relax—to assess their effects on ESM3’s performance. Our findings reveal that the use of a consistent identical structure, regardless of whether it is relaxed or variant, consistently enhances model performance across various datasets. This improvement is striking in few-show learning. However, performance deteriorates when different relaxed mutant structures are used for each variant. Based on these results, we advise caution when integrating distinct mutant structures into ESM3 and similar models.This study highlights the critical need for careful consideration of structural inputs in protein binding affinity prediction.

## 1 Introduction

The global COVID-19 pandemic has highlighted the critical role of protein-protein interactions in virus behavior, particularly through mutations such as amino acid substitutions. These mutations can disrupt existing interactions and alter binding energies ΔΔ*G* = Δ*G*_*MUT*_ *−* Δ*G*_*WT*_, impacting the structures and functions of proteins. The understanding of these dynamics, especially the interactions between the receptor binding domain (RBD) and antibodies or Angiotensin-converting enzyme 2 (ACE2), is crucial due to their implications in viral infectivity and evasion mechanisms [1].

Traditional experimental methods, including high-throughput experiments like deep mutational scan [2], while reliable, are often limited by their time-intensive and costly nature, making them impractical in urgent pandemic responses. This has led to an increased reliance on computational methods, which have shown to be invaluable not only in identifying potential viral threats but also in facilitating the rapid development and optimization of therapeutic antibodies [3] [4].

Computational approaches to studying protein dynamics can be broadly categorized into traditional biophysical methods and machine learning or deep learning techniques. Traditional biophysical methods, such as those utilizing force fields in tools like FoldX [5] and Rosetta [6], relying on empirical interatomic interactions to predict folding stability or binding affinity While effective, the highest accuracy in these methods often demands extensive computational resources, such as those required for Free Energy Perturbation [7].

Machine learning models and pLMs become a popular alternative to these resource-heavy physical simulations. Despite the inability to validate variant structures, the integration of 3D structural data into machine learning models is transforming the landscape of PPI prediction. Different models use different relaxation methods to obtain variant structure, in order to predict free energy difference ΔΔ*G* for protein-protein complexes. DDAffinity[8], a message passing neural network, uses FoldX [5] BuildMutant for both the wildtype and variant structures. DDGPred [9], a deep neural network, uses Rosetta Relax [10] to obtain mutated structures to predict. EGRAL[11], a graph neural network, uses Rosetta Relax and Backrub [12], in a pipeline close to the conformation sampling of flex ddg [6] method. It allows them to obtain variant structure for SKEMPI2.0 dataset [13] which only provides wildtype (wildtype) structures in order to predict mutational property changes. Unibind,[14] a Graph Neural Network, uses EvoEF2 BuildMutant [15] to obtain the structure graph.

While these papers concentrate on refining the architecture of their models to enhance performance on pretraining tasks and subtasks, comparatively little attention is given to evaluating the impact of these structural acquisition pipelines. These set the stage for our investigation into the ESM3 model [16], a multimodal pLM that, unlike previous ESM models [17] which is trained on sequence only, natively incorporates sequence data, all-atom coordinates, and functional annotations. This capability renders ESM3 exceptionally versatile, allowing it to generate embeddings from either sequence-only inputs or combined sequence and structure inputs. Importantly, there are no stringent requirements for the structures provided, nor is there a need for preprocessing these structures. This flexibility means the method to obtain protein structures can be modified easily and evaluated as part of the end-to-end pipeline. In the following work, we demonstrate that (i) identical protein structures significantly enhances the binding affinity predictions of ESM3, and (ii) the performance of the model is degraded by using a relaxed structure for each variant, compared to using the one identical structure of any variant.

## 2 Methodology

### 2.1 ESM3 and model inputs

The model we benchmarked is composed of ESM3 and a regression head. We have extracted the hidden representation used by output heads after the attention mechanism. Additionally, it is crucial to highlight that the geometric encoder is invariant to rotations and translations, so that alignment is not needed. ESM weights are kept frozen. The embeddings are mean-pooled to obtain a complex embedding, instead of a tensor of residue embeddings. We renormalized the embeddings and binding affinities ΔΔ*G* by mean and variance. In our results, we report Spearman correlations on several high-throughput datasets: the Bloom dataset[2], which consists of a deep mutational scan of the SARS-CoV-2 RBD binding to ACE2, covering 3,803 single-mutation variants of the RBD. The Desai dataset [18] [19], which includes 2^15^ = 32, 768 combinatorial mutational data of RBD binding with ACE2 and LY-CoV555, CB6, REGN10987 and S309 antibodies. It is crucial to note that for these datasets, all mutations are on the RBD.

Our comparison identifies the SVM, implemented with default settings in scikit-learn [20], as the most effective regression head for these datasets, as detailed in SI Figure 7. Consequently, we will utilize the SVM for subsequent analyses. However, it is important to note that our findings should not depend on the choice of the regression model.

We evaluate three distinct types of inputs: sequence-only, sequence paired with identical structure, and sequence paired with variant structures. For sequence and identical structure analyses, we use the ProteinChain class from ESM3 repository to load the desired PDB file, which reads both the sequence and atom coordinates. We can then modify the returned object to change the sequence while keeping structure preserved. For sequence and variant structures, we have the flexibility to directly load the PDB file or adjust the sequence to match the desired variant, depending on whether the sequence aligns with the target variant.

### 2.2 Acquiring variants structures

Computational techniques such as in silico mutations, rotamer optimization, and energy minimization are typically used to generate variant structures. These methods are based on the physical principle that the most stable structures are also the most probable, due to the conformation space following a Boltzmann distribution. At low temperatures, the minimum energy state serves as an approximation for the minimum of free energy, allowing for the representation of a protein by a single structure.

In line with these principles, our initial step involves performing an in silico mutation using EvoEF2. The EvoEF2 BuildMutant commandt[15], which performs rotamer optimization to mutate specific residues using pre-defined libraries of side chain conformations. After mutation, there are two popular protocols: (i) Rosetta Relaxation protocol [10] is used to obtain variant structure with alternating side chain and backbone optimization. (ii) Gromacs energy minimization uses a steep descent energy minimization method with force field CHARMM36.[21].

In the category of sequence paired with variant structures, three widely used relaxation protocols for obtaining variant structures were evaluated: (i) EVoEF2 alone (ii) EvoEF2 with Rosetta Relax, and (iii) EvoEF2 with Gromacs energy minimization. The wildtype RBD structure is extracted from PDB file 6XF5[22], a trimer of the isolated spike protein. In the latter experiments, unless specifically mentioned, the training size is 100 random samples.

Additionally, as a control, we conduct Molecular Dynamics(MD) simulations using Gromacs and the CHARMM36 force field to generate structures under thermodynamic fluctuations for the wildtype. Structures are extracted every 2 femtoseconds and assigned to specific variants. Furthermore, we introduce noisy structure variants by applying Gaussian noise with a 0.1 Ångströms(Å) variance to all atoms in an EvoEF2-relaxed structure. This method helps us assess the impact of structural perturbations on model performance.

We used Intel Xeon Sapphire Rapids CPUs and A100 GPUs. Rosetta Relax required approximately 10 minutes per sample, while Gromacs took about 20 minutes per sample, each using a single CPU core. The MD simulation for the wildtype was completed in 24 hours using a single A100 GPU and 15 CPU cores. Meanwhile, ESM3 generated embeddings within an hour on a single GPU, without any inference optimization. Notably, the EvoEF2 BuildMutant was quite efficient, processing roughly 30,000 variants serially in just about one hour.

## 3 Results

Firstly, we evaluated the performance of ESM3 using only sequence data, as well as when paired with identical structures. As shown in Figure 1, while the performance of ESM1, ESM2 and ESM3 varies when sequence is the only input, the performance of ESM3 is consistently superior when the sequence is paired with an identical structure. This enhancement is also significant across various datasets we tested, as shown in SI Figure 12.

**Figure 1:**
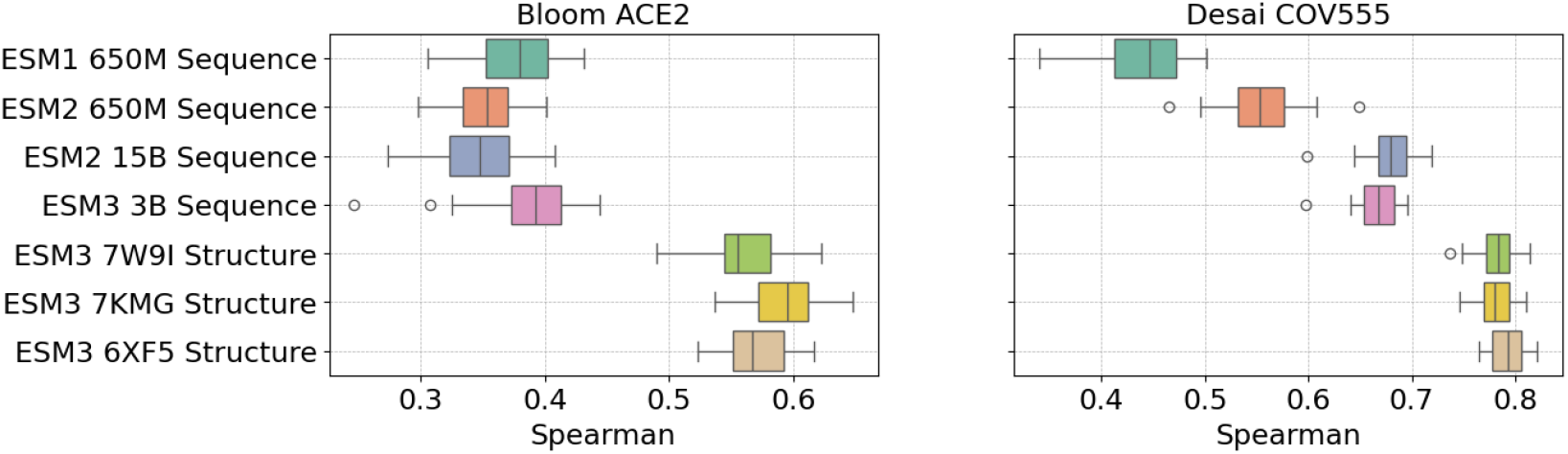
**Left:** Spearman correlation for Models: ESM1, ESM2, and ESM3 sequence only and sequence plus selected structures, evaluated on Bloom dataset for ACE2 binding affinity. 7W9I is the SARS-CoV-2 RBD bound to ACE2, 7KMG is RBD binding with the LY-CoV555 antibody, 6XF5 is the SARS-CoV-2 spike protein with RBD in the down position **Right:** Same analysis on the Desai dataset for RBD and LY-CoV555 antibody binding affinity.

One might anticipate that using RBD structures from RBD-ACE2 binding interactions would yield better results when predicting the binding between RBD mutants and ACE2. As there are minor structural change on the binding surface due to PPI. However, our findings show no significant difference in predictive accuracy. For instance, the structure of 7W9I [23], which is the SARS-CoV-2 RBD bound to ACE2, did not demonstrate enhanced predictive accuracy when predicting on Bloom RBD-ACE2 dataset. Similarly, 7KMG [24], which contains RBD binding with the LY-CoV555 antibody, and 6XF5 [25], the SARS-CoV-2 spike protein with RBD in the down position, also did not exhibit a significant difference in predictive performance in either task shown in Figure 1. We have also tested other PDB structures containing various variants and antibodies, from which we use only the RBD chain, as shown in SI Figure 6, and found no significant differences. This suggests that, although there are minor structural differences in these PDB structures, the ESM3 model could not effectively utilize these variations to enhance prediction accuracy.

Secondly, we evaluate the performance of ESM3 when it is provided with sequences and structures for each mutant, sourced from various pipelines. As indicated previously, using the wildtype structure—referred to here as ‘Wildtype’—enhances the performance of the ESM3 model compared to using sequence data alone.

Interestingly, as shown in Figure 2, utilizing the EvoEF2 generated mutant structure does not show any observable effects on the predictive power of ESM3 across all datasets we tested. Furthermore, after using EvoEF2 to introduce mutations, we applied two popular relaxation methods. Surprisingly, the minimized mutated structures produced via Rosetta or Gromacs exhibited lower Spearman correlations, aligning their performance with that of sequence-only inputs.

**Figure 2:**
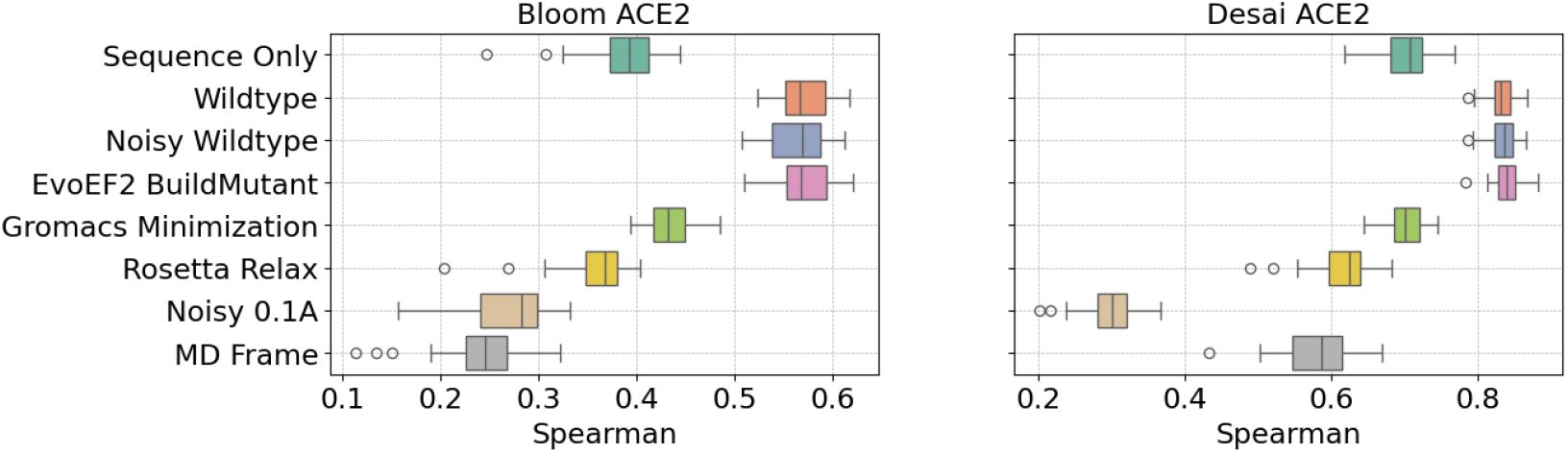
**Left:**Spearman correlation for sequence plus variant structures obtained through different structure acquisition pipelines, evaluated using ESM3 on the Bloom ACE2 **Right:** Same analysis on the Desai dataset for ACE2 binding affinity. Example of aligned strucures are shown in SI Figure 4 and 5

Consequently, we tested two additional structural pipelines: the ‘MD Frame’ in which each mutant is provided with a structure extracted every 2 femtoseconds during MD simulations, and the ‘Noisy 0.1 Å’, which involves a Gaussian noise with mean 0 and variance 0.1 Å applied to all atoms in an EvoEF2-generated configuration. Interestingly, despite the minimal perturbation in the ‘MD Frame’ and ‘Noisy 0.1A’, there is a significant reduction in accuracy, even falling below that of sequence-only embeddings. This indicates that the embeddings are highly sensitive to noise introduced by processing pipelines. These results are consistent on other datasets we tested, all summarized in SI Figure 11. Further specifics for each dataset can be found in (LY-CoV555: 14, CB6: SI Figure 15, REGN10987: SI Figure 16, S309: SI Figure 17 and ACE2: SI Figure 13).

Lastly, Gromacs minimization and Rosetta Relax were run on the wildtype structure and provided as structure to ESM3. These embeddings with the same structure gave all equal performance with Wildtype pipeline, as shown with “Noisy Wildtype” in Figure 2 and in SI Figure 13-17. This effectively suggests that small perturbation is not an issue provided that the same structure is used for all variants. Hence, ESM3 seems to be sensitive to RMSD when provided with protein structure variants. It is important to note that the structural variations induced by these pipelines are extremely subtle. To accurately measure the differences between the structures generated by these pipelines, we analyzed the Root Mean Square Deviation (RMSD) distribution by comparing the structures associated with the wildtype variant to those relaxed using the same method. Across all structures, the RMSD consistently remains below 2Å, as shown in Figure 3 which is still within the limits of PDB measurement resolution.

**Figure 3:**
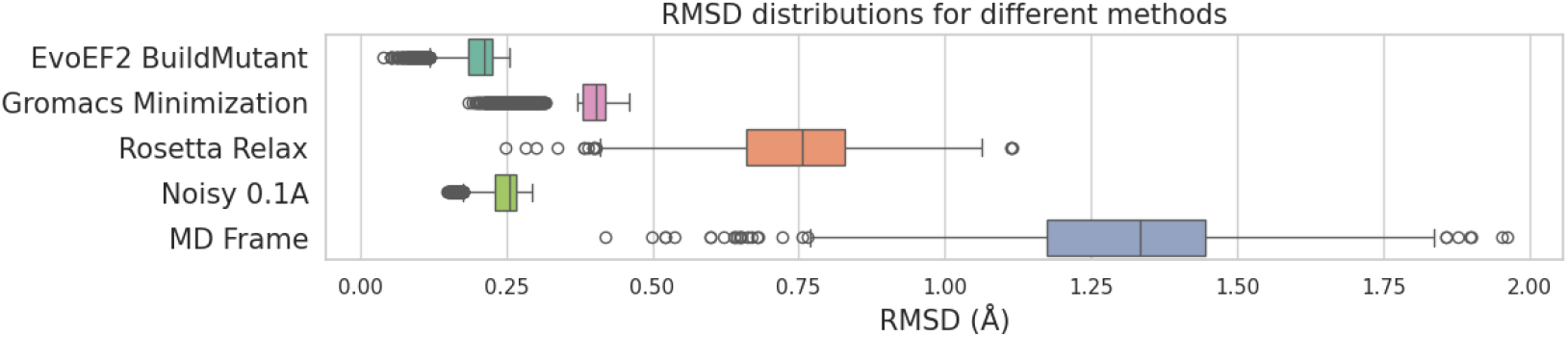
RMSD distribution for structures acquired using different methods.

Nonetheless, we can observe that the ranking of methods by mean RMSD—Minimized Gromacs, Relax Rosetta, and MD Frame—correlates inversely with their Spearman correlation rankings, discounting the noisy structure, which is unphysical. The MD Frame, showing the highest RMSD differences, performed the worst, whole the Gromacs Minimization performed the best with the lowest RMSD. This is counter-intuitive to the physical understanding that under thermodynamic fluctuations, MD and relaxed structures should represent the actual protein structure and thus be treated the same by the model.

### 3.1 Embeddings similarities

A further analysis of the embeddings before the renormalization shows a very high similarity of the embeddings whose structures are generated from the same pipeline. The cosine similarity of these embeddings compared to its own wildtype embedding is at least 0.99, as shown in Table 1. This indicates that the generated embeddings is contained in a very small subspace of ESM3 hidden representation space. This may explain the sensitivity of ESM3 given minimal change on the variant structures: it probably reaches the accuracy limit of this model pretrained on a very diverse dataset of proteins.

## 4 Discussions

Various studies utilize mutant structures generated by computational methods as inputs to their models for predicting PPI. In our work, we aim to benchmark these pipelines. We have adapted ESM3 for use as an encoder for PPI prediction. This adaptation was tested across several protein binding datasets with various mutant structures and different structure minimization pipelines. Initially, we observed that although using identical PDB structures enhances the model’s performance in predicting variant binding affinities, using identical RBD structures from different RBD-protein complexes does not significantly impact results. This finding supports the use of identical structures as a priori information in ESM3 for PPI prediction, even if these structures do not perfectly represent the actual protein configuration during binding.

Additionally, providing ESM3 with mutant structures generated by EVoEF2 instead of the structure does not lead to an increase in model accuracy. Furthermore, we noted that ESM3 exhibits sensitivity when provided with structural variations, whether they arise from relaxed structures, thermal noise in MD simulations, or artificially introduced Gaussian noise. To mitigate such sensitivity, constraints could be applied, for example, on amino acids far from mutation sites. This would allow for exploration of relevant parts of the conformation space while limiting displacement in the unaffected regions of the protein, leading to a smaller RMSD.

Our findings highlight the critical need for enhanced computational pipelines that better accommodate variants with measurable structural changes, given that none of the mutant-generating pipelines we tested improved model performance. Additionally, our results highlight the importance of further evaluating the effects of mutant structure pipelines on predicting PPIs and other task, both in pLMs and other models that require structural inputs.

Based on our findings, researchers using ESM3 to predict mutant PPIs might consider the following: (i) Researchers could potentially bypass MD relaxation or energy minimization, opting to use the PDB file directly from the PDB databank.(ii) Caution should be exercised when using relaxed structures generated by EvoEF2. Furthermore, while not tested in this paper, we encourage the evaluation of the impact of relaxation for pretrained or end-to-end models, to ensure that the use of the relaxation pipelines effectively improves the performance of the model while not overfitting the generated mutant structures.

## 5 Code Availability

The code for reproducing our experiments is available at https://github.com/Dianzhuo-Wang/esm3-structural-inputs

