## Supplementary material for "More Structure, Less Accuracy: ESM3’s Binding Prediction Paradox": SI

### A Appendix

#### A.1 Similarity of embeddings

| Method | Mean cosine similarity | Mean Euclidean distance |
|---|---|---|
| Wildtype | $0.99994 \pm 2.2 \times 10^{-5}$ | $96.7 \pm 21.3$ |
| Relax Rosetta | $0.99963 \pm 9.35 \times 10^{-5}$ | $257 \pm 35$ |
| Minimized Gromacs | $0.99962 \pm 1.15 \times 10^{-4}$ | $260 \pm 40$ |
| MD Frame | $0.99933 \pm 1.40 \times 10^{-4}$ | $350 \pm 38$ |
| Noisy 0.1Å | $0.99976 \pm 9.51 \times 10^{-5}$ | $213 \pm 44$ |

Table 1: Comparison of methods based on cosine similarity and Euclidean distance

#### A.2 Aligned PDB structures

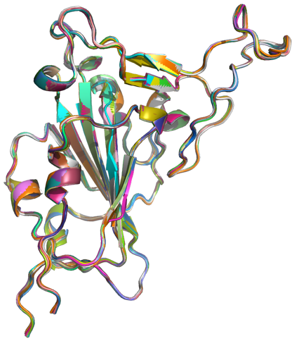

Figure 4: 40 randomly selected RBD structures from the Bloom dataset, obtained via the "MD Frame" pipeline.

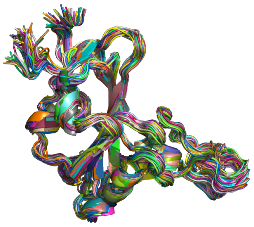

Figure 5: 40 randomly selected RBD structures from the Bloom dataset, obtained via the "Gromacs Minimization" pipeline.

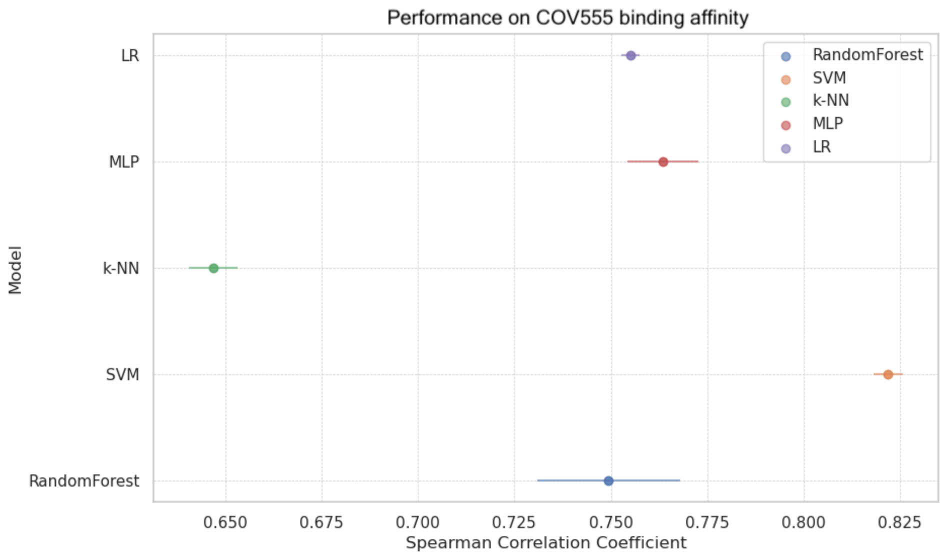


Figure 7: Choice of regression head for RBD binding affinities with COV555 antibody on Desai Dataset

#### A.3 RBD structures extracted from PDB file performan similarly

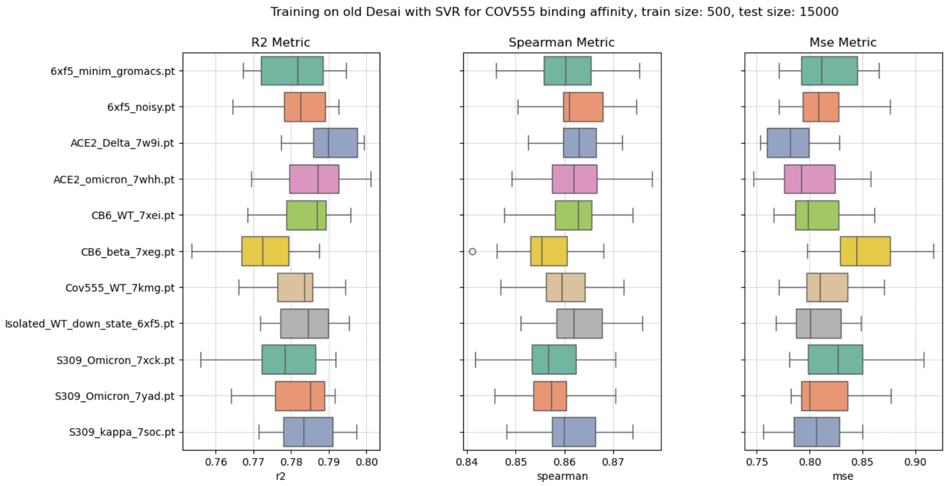


Figure 6: Performance for binding affinity with COV555 for RBD structures extracted from different PDB files

#### A.4 Choice of regression head

A comparison over all antibodies binding affinities in Desai Dataset suggest that SVM regressor performs the best.

##### A.5 ESM1 vs ESM2 vs ESM3

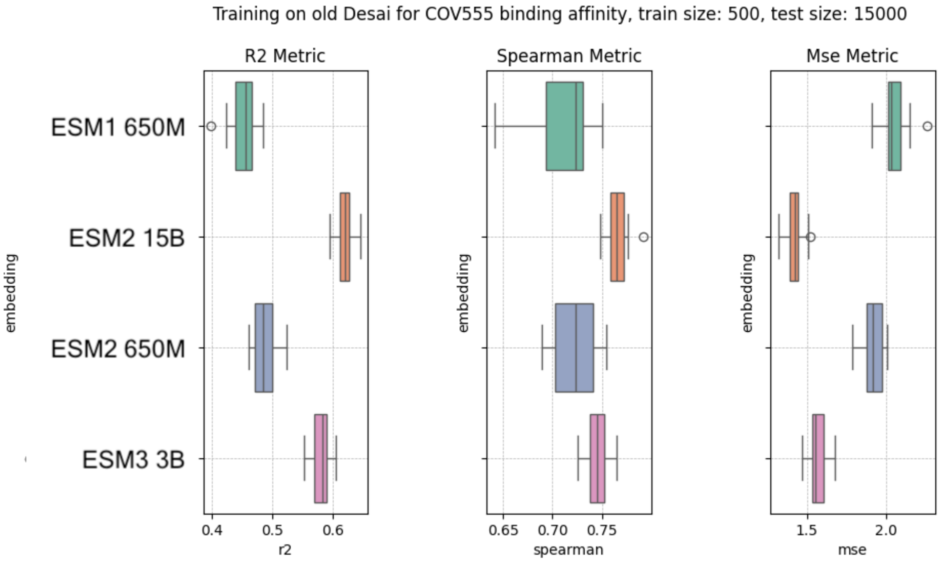


Figure 8: Performance of sequence-only for ESM1 650m, ESM2 650m, ESM2 15b, ESM3

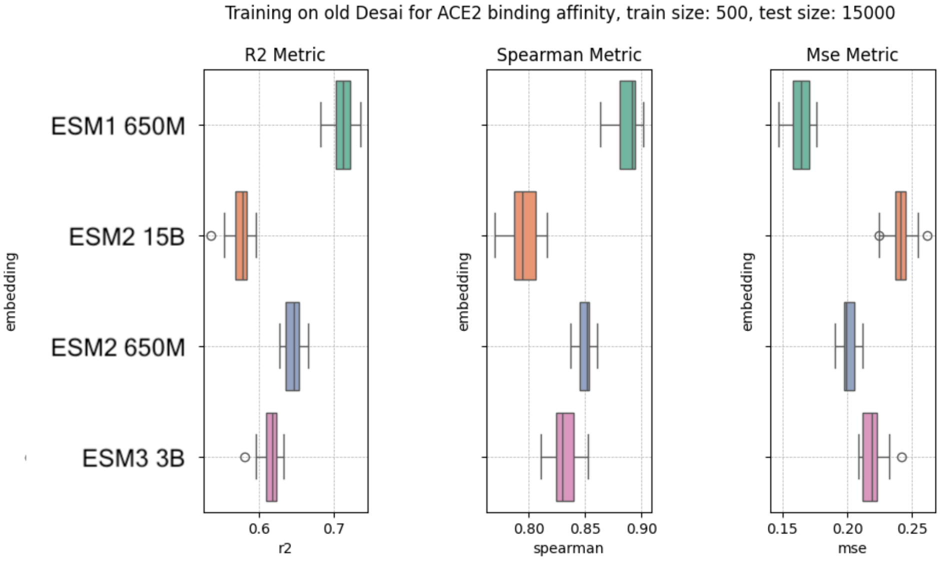


Figure 9: Spearman Correlation for RBD binding affinity with ACE2

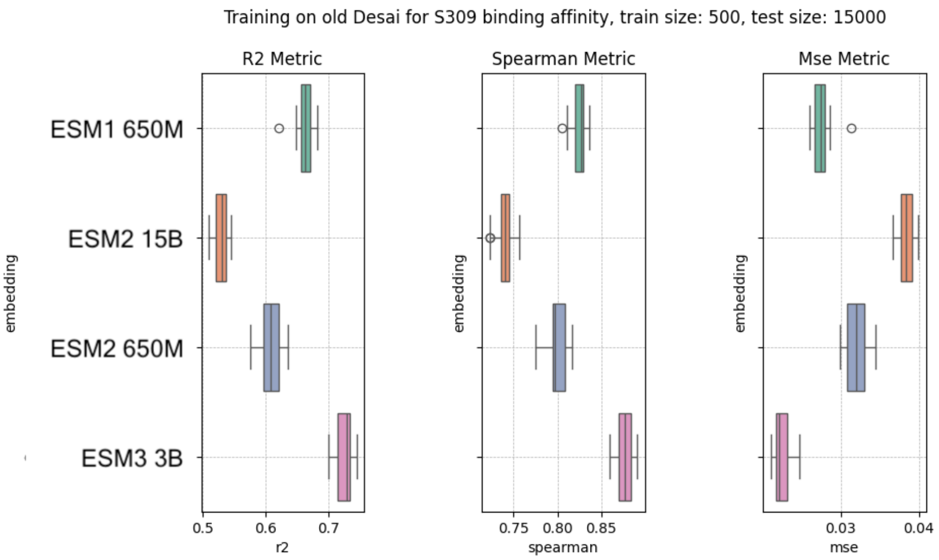


Figure 10: Spearman Correlation for RBD binding affinity with ACE2

##### A.6 Performance on binding affinity prediction with antibodies on Desai Dataset

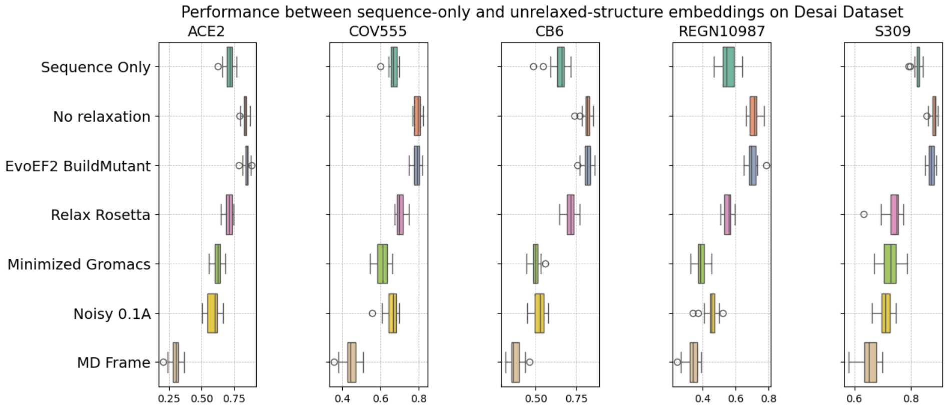


Figure 11: Spearman Correlation for RBD binding affinity with other proteins. Training size is 100 random samples.

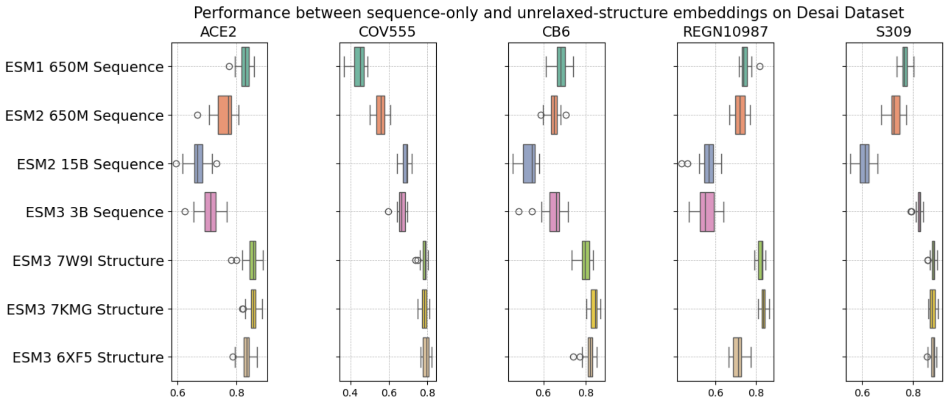


Figure 12: Spearman Correlation for RBD binding affinity with other proteins comparing sequence-only embeddings and sequence and structure. Training size is 100 random samples.

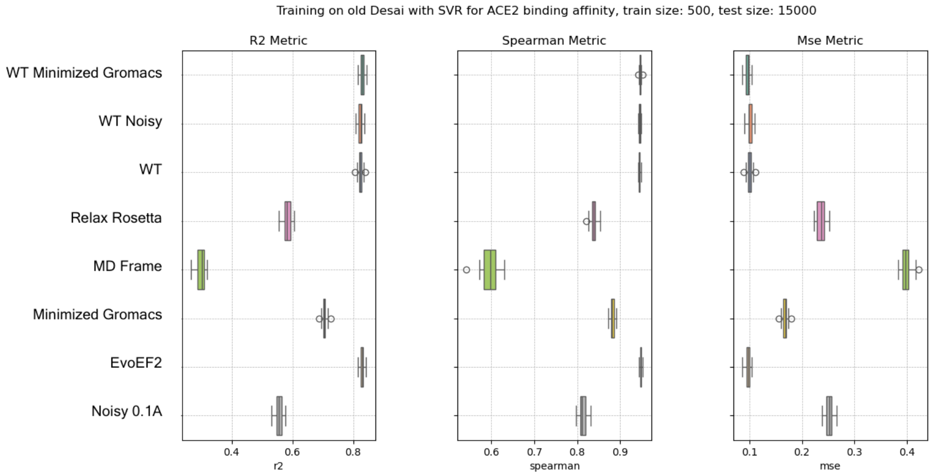


Figure 13: Spearman Correlation for RBD binding affinity with ACE2

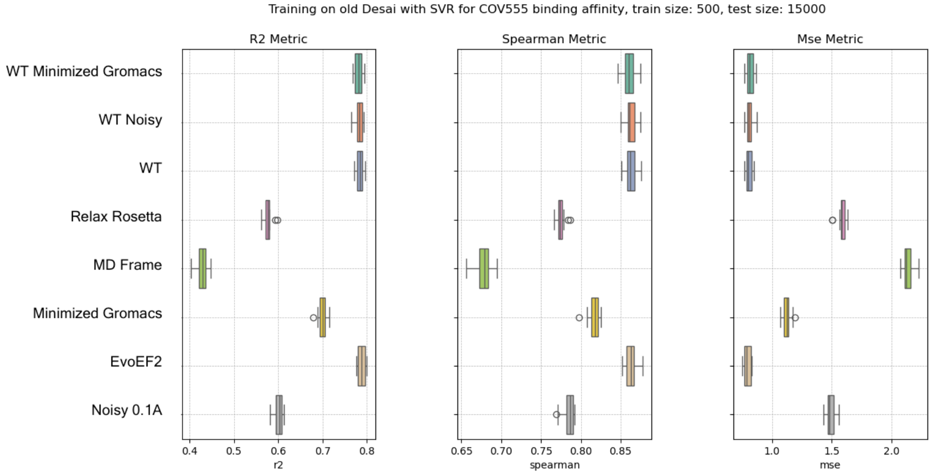


Figure 14: Spearman Correlation for RBD binding affinity with COV555

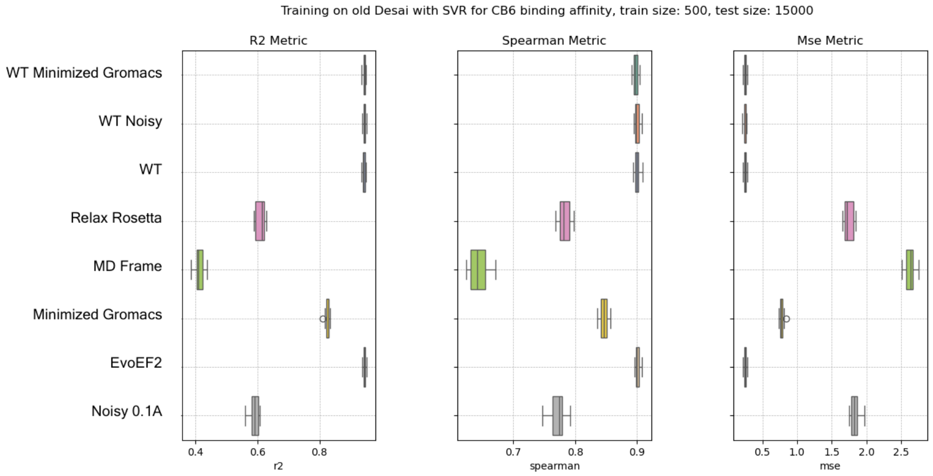


Figure 15: Spearman Correlation for RBD binding affinity with CB6

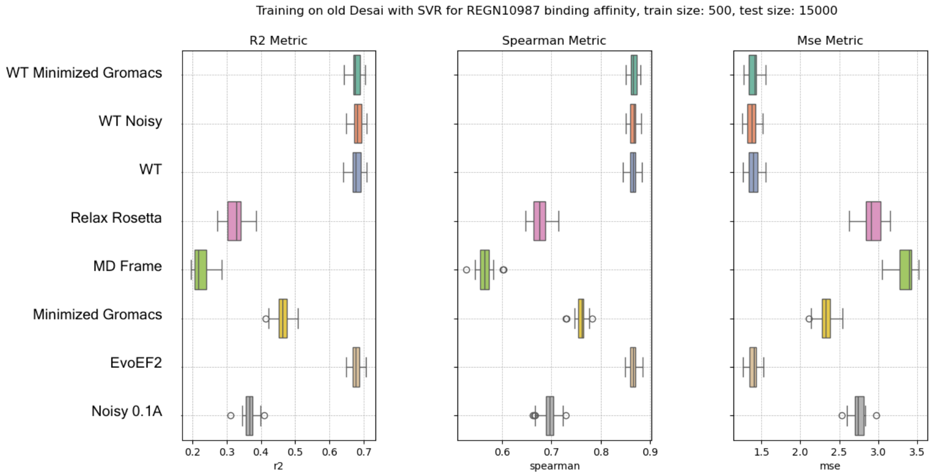


Figure 16: Spearman Correlation for RBD binding affinity with REGN10987

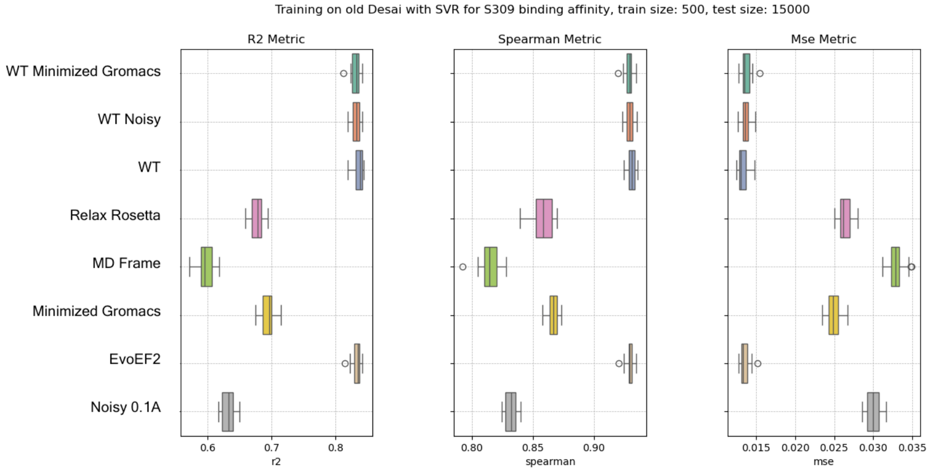


Figure 17: Spearman Correlation for RBD binding affinity with S309
